# Computational anatomy and geometric shape analysis enables analysis of complex craniofacial phenotypes in zebrafish

**DOI:** 10.1101/2021.02.12.431035

**Authors:** Kelly M. Diamond, Sara M. Rolfe, Ronald Y. Kwon, A. Murat Maga

**Affiliations:** Center for Developmental Biology and Regenerative Medicine, Seattle Children’s Research Institute, Seattle, WA, USA; Friday Harbor Marine Laboratories, University of Washington, San Juan, WA, USA; Department of Orthopedics and Sports Medicine, University of Washington, Seattle, WA, USA; Institute for Stem Cell and Regenerative Medicine, University of Washington, Seattle, WA, USA; Division of Craniofacial Medicine, Department of Pediatrics, University of Washington, Seattle, WA, USA

**Keywords:** cranial morphology, osteogenesis imperfecta, geometric morphometrics, computational anatomy

## Abstract

Due to the complexity of fish skulls, previous attempts to classify craniofacial phenotypes have relied on qualitative features or 2D landmarks. In this work we aim to identify and quantify differences in 3D craniofacial phenotypes in adult zebrafish mutants. We first estimate a synthetic ‘normative’ zebrafish template using microCT scans from a sample pool of wildtype animals using the Advanced Normalization Tools (ANTs). We apply a computational anatomy (CA) approach to quantify the phenotype of zebrafish with disruptions in *bmp1a*, a gene implicated in later skeletal development and whose human ortholog when disrupted is associated with Osteogenesis Imperfecta. Compared to controls, the *bmp1a* fish have larger otoliths and exhibit shape differences concentrated around the operculum, anterior frontal, and posterior parietal bones. Moreover, *bmp1a* fish differ in the degree of asymmetry. Our CA approach offers a potential pipeline for high throughput screening of complex fish craniofacial phenotypes, especially those of zebrafish which are an important model system for testing genome to phenome relationships in the study of development, evolution, and human diseases.

**Summary statement:** A computational anatomy approach offers a potential pipeline for high throughput screening of complex zebrafish craniofacial phenotypes, an important model system for the study of development, evolution, and human diseases.

## Introduction

The fish craniofacial skeleton has long functioned as an archetype system for elucidating genetic and environmental contribution to phenotype in vertebrates. In the context of development, studies have focused on the genetic mechanisms that shape the cranial skeleton (Kimmel et al., 2020; Miller et al., 2007). Craniofacial analyses have been used to understand the pathways that have enabled morphological evolution (Kimmel et al., 2005), phenotypic plasticity (Navon et al., 2020), and adaptive radiations (Powder and Albertson, 2016) in fishes. Additionally, zebrafish are developing as a model system for quantifying phenotypic variability associated with human bone diseases, such as Osteogenesis Imperfecta (Busse et al., 2019; Gistelinck et al., 2018; Kwon et al., 2019). A longstanding challenge to analyzing the fish craniofacial skeleton is accurately capturing phenotypes that involve subtle alterations and complex 3D changes, including potential asymmetric alterations.

The traditional methods for quantifying cranial morphology use manually-placed homologous landmark points on 2D images of the lateral view of the head (i.e. Sidlauskas, 2008). However, manual placement limits potential for rapid-throughput applications. Further the requirement for homologous structures limits landmark placement across the skull, and hence may miss the phenotypic variation in these areas. While microCT can help realize 3D structures, 3D landmark placement is complex as visualizations are dependent on both the scanner and rendering software settings used. Moreover, because of the close proximity of bones, segmentation-based approaches that are useful for axial skeleton are not amenable to those in the head. There is an urgent need to develop robust methods for phenotyping in the craniofacial skeleton that are sensitive to complex 3D changes while being amenable to rapid-throughput analyses.

Here, we propose using an atlas-based computational anatomy (CA) approach to build a template and then using a pseudo-landmark pipeline to identify areas of the skull that vary among mutant and wildtype fish. Atlas-based approaches estimate an unbiased anatomical ‘template’ from a group of images (Guimond et al., 2006), and then use this template to the basis to assess shape differences among groups of interest (Ashburner and Friston, 2000). Atlas-based approaches have been used to characterize phenotypes in many neuroimaging studies in humans, in fetal mice (KOMP2 project) as well as in the mouse cranial skeleton (Maga et al., 2017; Toussaint et al., 2020). We define pseudo-landmarks here as landmarks that are not homologous but evenly cover the surface of our fish skulls.

We apply these methods to zebrafish with mutations in *bmp1a*, a gene implicated in later skeletal development. In humans, Bone Morphogenetic Protein 1 (BMP1) encodes for a secreted protein involved in procollagen processing. Individuals with mutations in BMP1 exhibit increased bone mineral density and recurrent fractures characteristic of Osteogenesis Imperfecta (OI; Asharani et al., 2012). Severe forms of OI are frequently associated with craniofacial abnormalities (Dagdeviren et al., 2019). Previous work in *bmp1a* and other zebrafish OI models have identified phenotypic abnormalities in the axial skeleton (Hur et al., 2017). However, due to the complicated structure of the fish cranial skeleton, craniofacial abnormalities in zebrafish OI models have mostly focused on qualitative phenotypes (Gistelinck et al., 2018), and little work has been done to quantify complex cranial phenotype. Here, we report complex craniofacial phenotype arising from disruptions in *bmp1a*. Our methods aim to quantify the cranial phenotype associated with mutations in zebrafish with minimal user intervention so that large scale studies can examine phenotype-genotype associations in the skeletal system in a high-throughput

## Methods

Generation of mutant animals and microCT scanning were described in Watson et al., 2020. We used a total of 23 wildtype fish from two clutches (“wildtype fish”) to build our atlas and used 12 *bmp1a* somatic mutants (“*bmp1a* fish”), from a single clutch. Watson et al., 2020 performed a comparison of *bmp1a* somatic and germline mutants and showed that somatic *bmp1a* mutants recapitulate germline *bmp1a* mutant phenotypes but possess additional phenotypic variability due to mosaicism. We focused our analyses on *bmp1a* somatic mutants as they provide a real-world sample of phenotypic variability likely to be encountered in CRISPR-based reverse genetic screens (Shah et al., 2015; Watson et al., 2020). Whole-body microCT images were acquired with a 21 micron voxel size.

### Atlas Building

To investigate potential asymmetry patterns, we built a symmetrical atlas of wildtype fish (N=23) by first reflecting all volumes along the sagittal plane. A symmetric atlas was generated using the antsMultivariateTemplateConstruction2.sh script as provided by the Advance Normalization Tools (Avants et al., 2014), using the default settings. Atlas building script initiates with a linear average of all samples, to which all samples are deformable registered to (Figure 1). The resultant deformation fields are applied to samples, and a new average is estimated and then used as a new reference for the next step of registrations. Four iterations were sufficient to obtain a symmetrical and anatomically detailed template.

**Figure 1.**
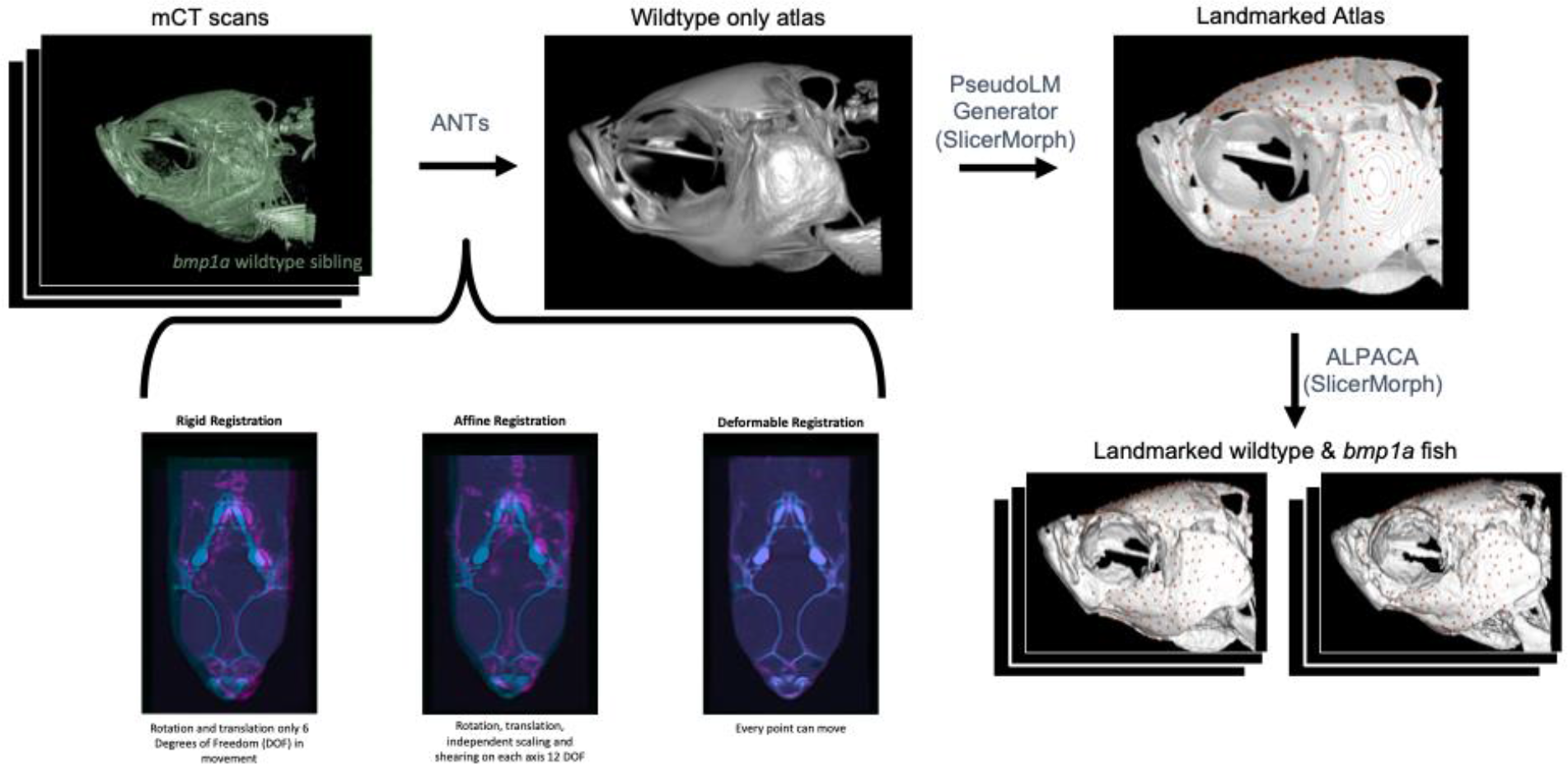
Pipeline for atlas building, pseudo-landmark generation, and transferring pseudo-landmarks to individual fish. Starting with microCT scans of wildtype fish ANTs, uses a series of rigid, affine, and deformable registrations to create an average image, or Atlas. The PseudoLMGenerator tool in SlicerMorph was used to place 372 pseudo-landmarks on the atlas. The ALPACA tool in SlicerMorph was used to transfer points from the atlas to wildtype and bmp1a fish for comparisons between groups.

### Atlas validation

First, we reviewed the resultant atlas qualitatively by investigating the spatial arrangement of bones in 3D rendering. To quantitatively validate the atlas and our computational anatomy framework, we created segmentations of individual otoliths from every sample manually using the open-source 3D Slicer program (Fedorov et al., 2012). We chose otoliths because they are dense, spread out along the dorsoventral axis of the crania, and do not touch any bones, which minimized the potential for error in our manual segmentations that serve as the ground truth data. The otoliths from the atlas were segmented in the same manner. Next, we deformably registered every sample, including the mutants, to our atlas using the ANTsR package, and applied the resultant transformation field to our atlas otolith segmentation to map them directly onto the subject space. From this mapping, we calculated the volumes of CA derived segmentation and statistically compared them to ground truth. All statistical analysis and image registrations were done using the R extensions of the ANTs ecosystem (Avants, 2020).

### Analysis of ZF cranial shape difference in wild-types and mutants

We first sparsely placed manual landmarks (N=8) on all subjects in our study and performed a Euclidian Distance Matrix Analysis (EDMA) on the manual landmarks using the EDMAinR package in R (https://github.com/psolymos/EDMAinR). In the EDMA analysis we used the *bmp1a* fish as the numerator and wildtype fish as the denominator. Landmarks used included (1) posterior most point of parietal (2) anterior most point of frontal (3) posterior most point of maxilla (4) left ventral most point of lower jaw (5) anteriodorsal most point of 1st vertebrae (6) right ventral most point of lower jaw (7) left postocular process and (8) right postocular process (Figure S1).

To examine the overall shape variation, we compared densely spaced pseudo-landmark points between *bmp1a* and wildtype fish. To place pseudo-landmark points on each of our specimens, we first created 3D models of from our ct volumes using the Segment Editor module of 3D Slicer (Fedorov et al., 2012). To generate a set of pseudo-landmark points on our atlas model, we used the PseudoLMGenerator module in the SlicerMorph extension of 3D Slicer which uses the original mesh geometry and a sagittal plane as the axis of symmetry, to generate a dense point cloud (Rolfe et al., 2021). One author (KMD) then went through the pseudo-landmarks and removed points that were on both jaws and the pectoral girdle using the MarkupEditor tool in 3D Slicer (Rolfe et al., 2021; Figure 1). Both of these structures are highly prone to plastic post-mortem deformation due to handling and preservation, as such they represent confounding non-biological variation. To transfer the pseudo-landmark points from the atlas to all other models in the study, we used the ALPACA module in the SlicerMorph extension of 3D Slicer, which uses linear and deformable point cloud registration (Porto et al., 2020). We used the default settings and skipped the scaling option to transfer pseudo-landmarks from the atlas to all meshes in our sample (Figures 1, S2).

To examine differences between *bmp1a* and wildtype fish, we ran a Generalized Procrustes Analyses (GPA) on the 372 pseudo-landmark points (pLMs), allowing all pLMs to slide along the surface, using the geomorph package in R (Adams and Otárola-Castillo, 2013). We ran a symmetry analysis on the GPA coordinates using the bilat.symmetry function in the geomorph package in R (Adams and Otárola-Castillo, 2013). From this output, we ran Procrustes ANOVAs to determine if the symmetric, and fluctuating asymmetric components of shape variation differ between groups. We also ran separate principal components analyses on both the symmetric and asymmetric components of variation from the symmetry analysis using the geomprph package in R (Adams and Otárola-Castillo, 2013). Visualizations were created in the SlicerMorph extension of 3D Slicer (Rolfe et al., 2021) and using ggplot in R (Wickham, 2016).

## Results & Discussion

When analyzing otoliths, we did not find significant differences between manually segmented volumes and atlas segmented volumes (t=-0.912, p=0.363; Figure S3), though there were some differences between some of the individual otoliths (Table S1; Figure S3). The most apparent difference between *bmp1a* and wildtype fish is that *bmp1a* fish have larger otoliths than wildtype fish, especially for the asteriscus, the largest otoliths in the zebrafish. This difference was consistent in both manually and CA segmented otoliths (Table 1; Figure S3). In contrast to bone formation, in which the mineral phase is primarily hydroxyapatite, otoliths are formed via an accumulation of calcium carbonate in the acellular endolymph of the fish inner ear (Payan et al., 2004). Previous work found higher tissue mineral density in *bmp1a* fish across the axial skeleton (Hur et al., 2017) and this result suggests potential influence of *bmp1a* on other pathways associated with mineralized tissues.

**Table 1.** Welch two sample t-test for difference between mutants and wildtype fish for each pair of manually segmented otolith volumes. We provide the mean volumes(x) for mutants and wildtype groups, degrees of freedom (df), test statistic (t), p value (p), and confidence interval (UCL-LCL).

| Otolith | x mutant<br>(mm<sup>3</sup>) | x wildtype<br>(mm<sup>3</sup>) | df | t | p | UCL | LCL |
| --- | --- | --- | --- | --- | --- | --- | --- |
| Left asteriscus | 0.044 | 0.039 | 15.975 | 3.383 | 0.004 | 0.002 | 0.008 |
| Right asteriscus | 0.044 | 0.040 | 14.62 | 3.232 | 0.006 | 0.002 | 0.007 |
| Left lapilus | 0.026 | 0.025 | 18.013 | 2.000 | 0.061 | -0.0007 | 0.003 |
| Right lapilus | 0.026 | 0.025 | 17.693 | 1.554 | 0.138 | -0.0004 | 0.003 |
| Left sagitta | 0.005 | 0.005 | 18.665 | 2.244 | 0.037 | 0.0003 | 0.009 |
| Right sagitta | 0.005 | 0.005 | 15.375 | 1.902 | 0.076 | -0.0005 | 0.008 |

In our EDMA analysis of the 8 manually placed landmark points, we found overall differences between *bmp1a* and wildtype fish (T=1.577, p<0.001). In comparing form distance ratios among all landmark pairs, we found less similarity between landmarks placed at the anterior portion of the head (landmarks 2-4 and 6-8) in our dataset compared to the two posterior most placed landmarks (landmarks 1,5; Figure S1). However, these landmarks were very sparsely placed and could be missing variation present in areas of the skull where traditional landmarks are sparse.

To see if there were areas of the skull that had greater variation than what could be determined from our EDMA analysis, we deployed a pseudo-landmark approach, placing 372 geometrically placed pseudo-landmarks across the outer surface of the cranial skeleton. In our symmetry analysis of pseudo-landmark points, we found significant differences in symmetry between groups for both the symmetric (F=3.573, Z=2.708, p=0.011) and asymmetric (F=3.830, Z=3.124, p=0.002) components of shape variation. The symmetric differences in shape variation between groups were concentrated in the anterior frontal bone and the dorsal portion of the operculum (Figure 2). While the asymmetric differences between groups were concentrated in the posterior portion of the parietal bone and ventral portion of the operculum (Figure 2).

**Figure 2.**
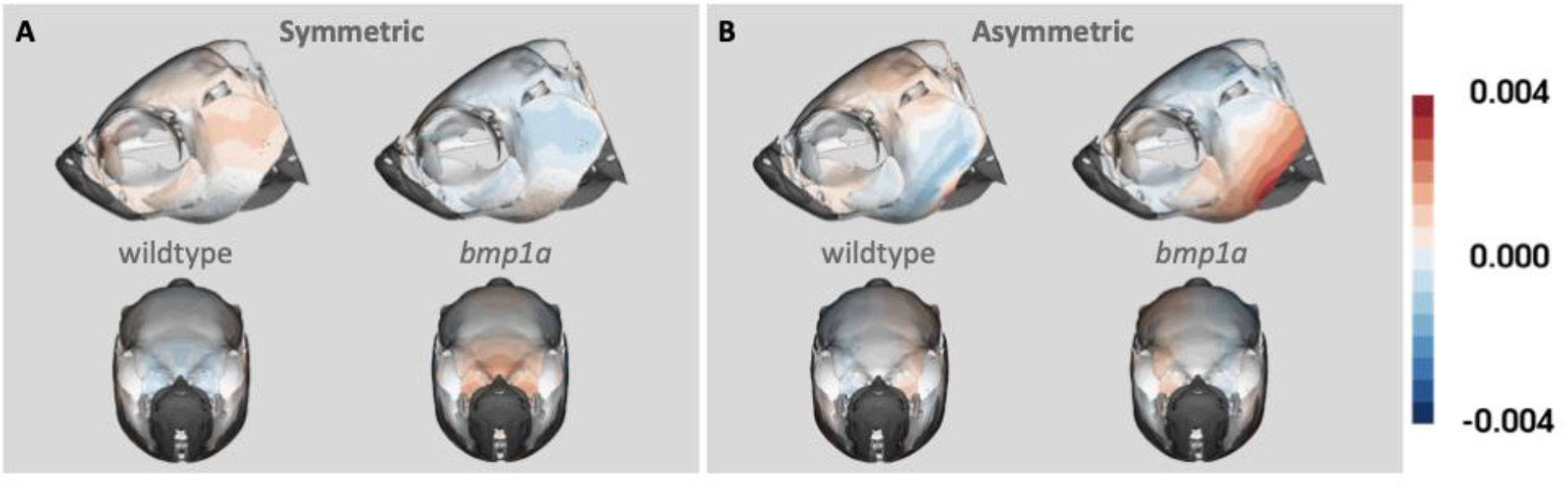
Heat map of (A) symmetric and (B) asymmetric components of shape variation. Lateral and anterior views are shown for each group (wildtype and *bmp1a*) within both components of shape variation. Colors show variation in shape from the symmetric atlas, with deeper colors representing greater variation from the atlas.

The results of separate PCA of each shape component suggest the asymmetric component of shape may be contributing more to the variation between groups in our dataset. For the symmetric component of variation, we found significant differences between *bmp1a* and wildtype fish along PC2, which explained 12.3% of the variation in the data (F=7.018; Z=2.006; p=0.002), but not along PC1, which explained 42.9% of the variation in the data (F=2.583; Z=1.124; p=0.092; Figure 3) or any other PCs. Whereas in the asymmetric shape space, we found differences between groups along PC1, which explained 35.0% of the variation, (F=6.305, Z=1.753, p= 0.009), but not along PC2, which explained 17.1% of the variation (F= 0.318, Z=-0.374, p=0.677; Figure 3), or any other PCs. When we visualize the first two principal components of the symmetry analysis, we find that the positive axis of the first principal component is influenced by the symmetrical and asymmetrical components of the posterior operculum (Figure 3). The negative axis of PC1 differs among the components of symmetry, with the symmetric component concentrated in the anterior portion of the frontal bone and the asymmetric component concentrated in the lateral parietal and supraocular regions (Figure 3). Very little variation is observed in the symmetric component of PC2, while the asymmetric component of this axis is again concentrated around the opercular and ocular regions (Figure 3). As we removed pseudo-landmark points associated with areas of the skull that varied due to preservation or scanning methods, these represent areas of interest for exploring how phenotype differs between mutant and wildtype fishes. Together these results provide evidence for phenotypic effects of the *bmp1a* mutation on the cranial phenotype of zebrafish. Future work should expand the number of families to ensure this is not unique to this particular family. We have shown how our pipeline can identify areas of greatest variation among groups of animals. In combination with additional morphological analyses, we hope this pipeline will enable researchers to better define the links between genotype and phenotype.

**Figure 3.**
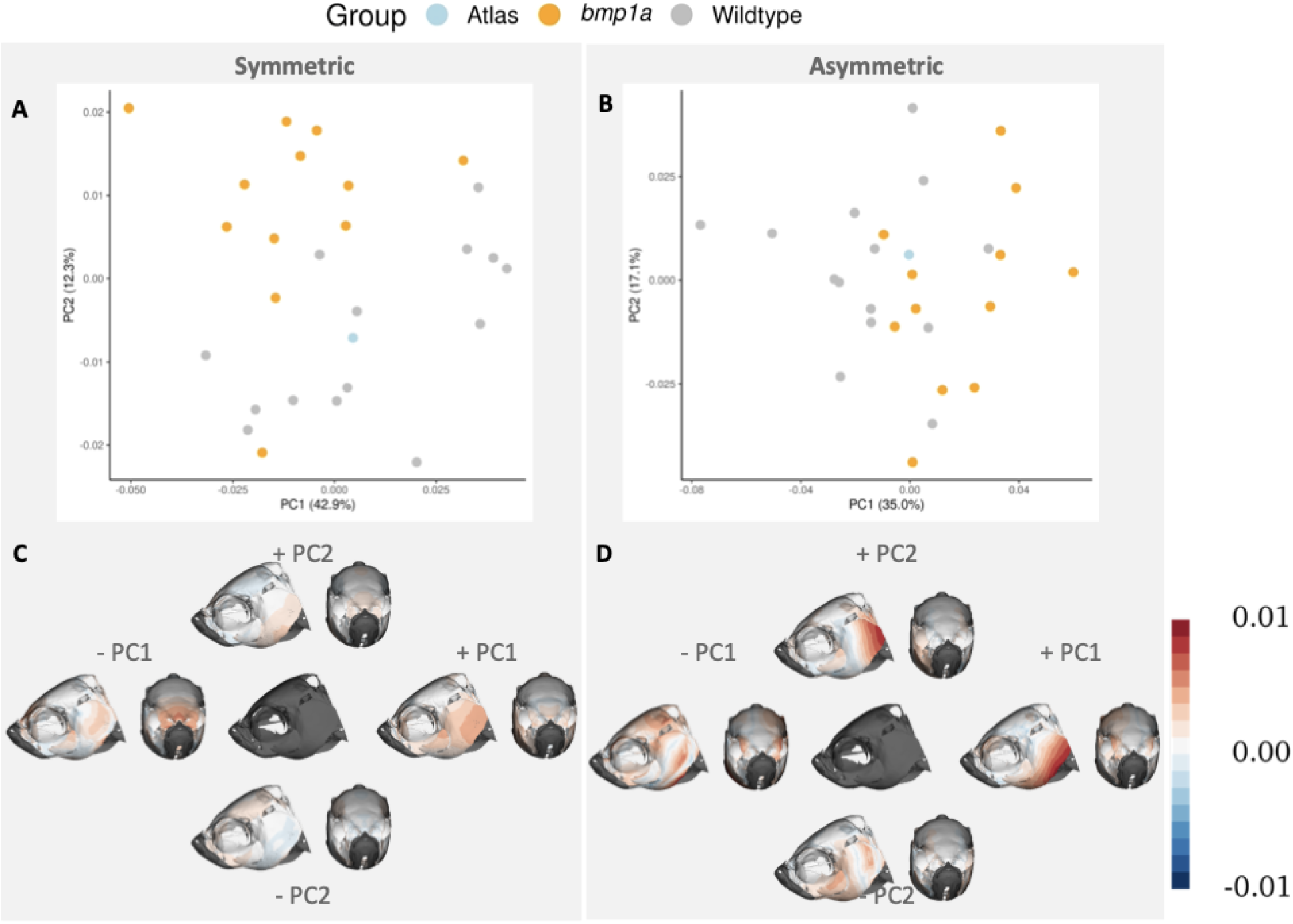
First two principal components of symmetry analysis. PC plots show separation of groups (represented by color) along the first and second PCs (A,B). Heat maps of the same PCs represent where shape variation occurs across each axis (C,D). Columns represent symmetric (A,C) and asymmetric (B,D) components of shape variation. The central image in C,D represent mean shape of each component. Color in C,D represents the Procrustes distance between the average shape and the shape occupying the ends of each PC axis.

## Acknowledgements

We thank members of the Maga lab and MSBL for feedback and development of this project. This project was partly supported by National Science Foundation grants An Integrated Platform for Retrieval, Visualization and Analysis of 3D Morphology from Digital Biological Collections (DBI/1759883) and Biology Guided Neural Networks for discovering phenotypic traits (OAC/1939505) to AMM. RYK was supported by NIH Grant AR074417.

## Author Contributions

Contribution of fish and microCT scans: RYK. Conceptualization and methodology: all authors, Writing: KMD; Editing and approval: all authors.

## Data availability

Atlas and pseudo-landmark points are available at: https://github.com/SlicerMorph/ZF_Skull_atlas/

