## Supplemental information for "Computational anatomy and geometric shape analysis enables analysis of complex craniofacial phenotypes in zebrafish"

**Table S1.** Welch two sample t-test for difference between manual and atlas-segmented otolith volumes for *bmp1a* and wildtype fish. We provide the mean volumes(x) for each method, degrees of freedom (df), test statistic (t), p value (p), and confidence interval (UCL-LCL).

| **Otolith** | **group** | **x atlas ($mm^3$)** | **x manual ($mm^3$)** | **df** | **t** | **p** | **UCL** | **LCL** |
|---|---|---|---|---|---|---|---|---|
| Left asteriscus | *bmp1a* | 0.042 | 0.044 | 21.971 | -0.902 | 0.377 | -0.006 | 0.044 |
| Left asteriscus | wildtype | 0.037 | 0.039 | 25.117 | -1.998 | 0.057 | -3.995 | 6.056 |
| Right asteriscus | *bmp1a* | 0.043 | 0.044 | 21.997 | -0.618 | 0.543 | -0.005 | 0.003 |
| Right asteriscus | wildtype | 0.037 | 0.040 | 20.891 | -2.744 | 0.012 | -0.005 | -0.0007 |
| Left lapilus | *bmp1a* | 0.024 | 0.026 | 22.000 | -3.074 | 0.006 | -0.005 | -0.0009 |
| Left lapilus | wildtype | 0.022 | 0.025 | 25.581 | -5.438 | <0.001 | -0.004 | -0.002 |
| Right lapilus | *bmp1a* | 0.023 | 0.026 | 20.94 | -2.490 | 0.021 | -0.005 | -0.0005 |
| Right lapilus | wildtype | 0.021 | 0.025 | 24.294 | -5.347 | <0.001 | -0.005 | -0.002 |
| Left sagitta | *bmp1a* | 0.006 | 0.005 | 17.619 | 1.998 | 0.061 | -0.00004 | -0.002 |
| Left sagitta | wildtype | 0.005 | 0.005 | 19.564 | 0.799 | 0.434 | -0.0003 | 0.0007 |
| Right sagitta | *bmp1a* | 0.006 | 0.005 | 16.603 | 1.934 | 0.070 | -0.0006 | 0.002 |
| Right sagitta | wildtype | 0.005 | 0.005 | 15.959 | 0.236 | 0.816 | -0.0005 | 0.0005 |

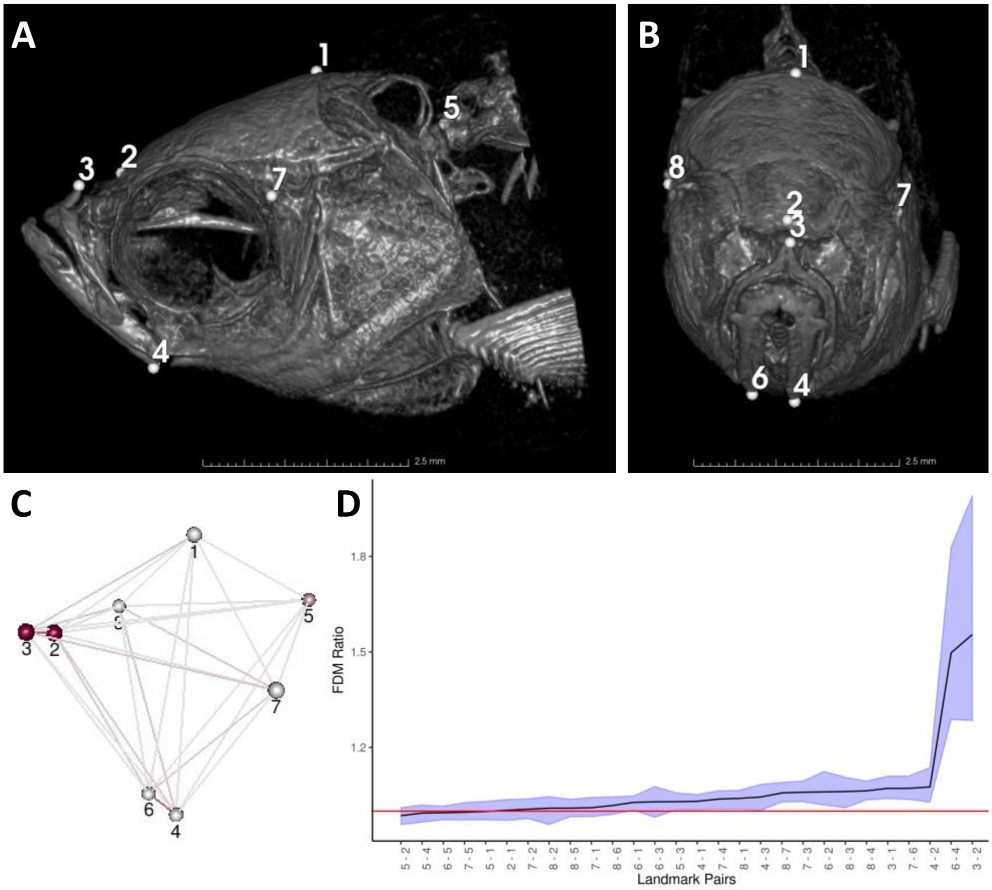


**Figure S1.** Euclidean Distance Matrix Analysis of manual landmarks. A) lateral and B) anterior views of example zebrafish ct scan with 8 landmark points used in analysis labeled, which include: (1) posterior most point of parietal (2) anterior most point of frontal (3) posterior most point of maxilla (4) left ventral most point of lower jaw (5) anteriodorsal most point of 1st vertebrae (6) right ventral most point of lower jaw (7) left postocular process and (8) right postocular process. C) Landmarks and Distances plotted with darker red coloration indicating more influential landmarks and distances. D) Form difference ratios for each set of landmark pairs (black line) and confidence intervals (shaded region). Individual values equal to 1 (red line) and confidence intervals that contain the value 1 indicate similarity between the samples for a given linear distance (Lele and Richtsmeier,1995).

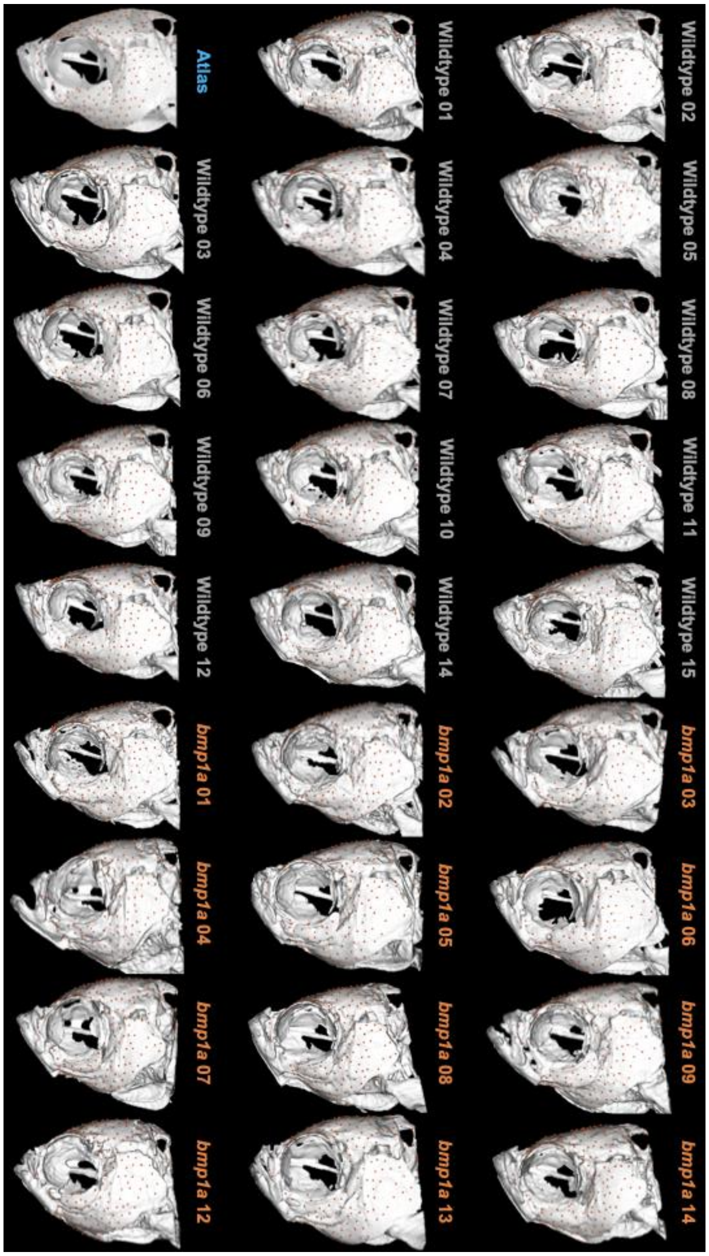


**Figure S2.** Left lateral view of fish meshes with pseudo-landmarks generated from the PseudoLMGenerator and transverse using the ALPACA modules in SlicerMorph. Name color represents groupings, the atlas is in blue, wildtype fish are in grey and *bmp1a* fish are in orange.

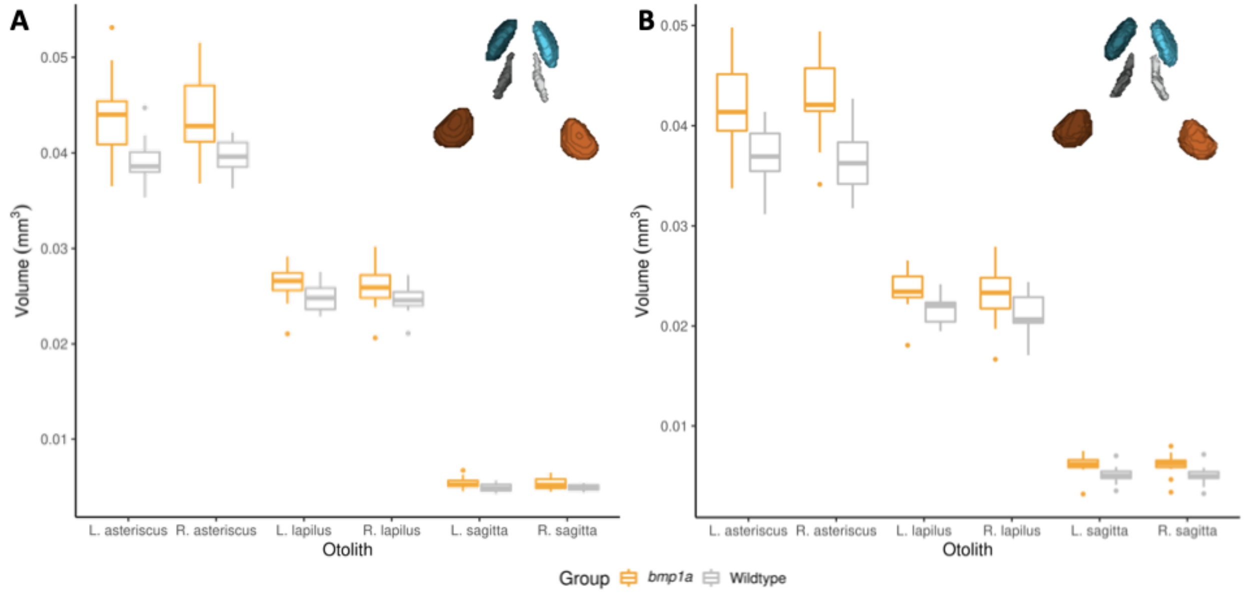


**Figure S3.** Boxplots of segmented otolith volumes. Wildtype (grey) and *bmp1a* mutant (orange) boxplots are shown for each of the six otoliths for both the (A) manually segmented volumes and (B) atlas segmented volumes. For each otolith, mutants have larger median volumes than wildtype fish (Table 1). Insets show the dorsal view of otolith segments (asterisk in blue, lapilus in orange, and sagitta in grey) with lighter colors on the left side of the head, and darker colors on the right.
